# A random forest model for predicting exosomal proteins using evolutionary information and motifs

**DOI:** 10.1101/2023.01.30.526378

**Authors:** Akanksha Arora, Sumeet Patiyal, Neelam Sharma, Naorem Leimarembi Devi, Dashleen Kaur, Gajendra P. S. Raghava

## Abstract

Identification of secretory proteins in body fluids is one of the key challenges in the development of non-invasive diagnostics. It has been shown in the part that a significant number of proteins are secreted by cells via exosomes called exosomal proteins. In this study, an attempt has been made to build a model that can predict exosomal proteins with high precision. All models are trained, tested, and evaluated on a non-redundant dataset comprising 2831 exosomal and 2831 non-exosomal proteins, where no two proteins have more than 40% similarity. Initially, the standard similarity-based method BLAST was used to predict exosomal proteins, which failed due to low-level similarity in the dataset. To overcome this challenge, machine learning based models have been developed using compositional features of proteins and achieved highest AUROC of 0.70. The performance of the ML-based models improved significantly to AUROC of 0.73 when evolutionary information in the form of PSSM profiles was used for building models. Our analysis indicates that exosomal proteins have wide range of motifs. In addition, it was observed that exosomal proteins contain different types of sequence-based motifs, which can be used for predicting exosomal proteins. Finally, a hybrid method has been developed that combines a motif-based approach and an ML-based model for predicting exosomal proteins, achieving a maximum AUROC 0.85 and MCC of 0.56 on an independent dataset. The hybrid model in this study performs better than the presently available methods when assessed on an independent dataset. A web server and a standalone software ExoProPred has been created for the scientific community to provide service, code, and data. (https://webs.iiitd.edu.in/raghava/exopropred/).

**Keypoints:**

- Exosomal proteins or non-classical secretory proteins are secreted by via exosomes
- A method has been developed for predicting exosomal proteins
- Models have been trained, tested, and evaluated on non-redundant dataset
- Wide range of sequence motifs have been discovered in exosomal proteins
- A web server and standalone software have been developed

## Introduction

Protein secretion is crucial for wide range of functions including communication among cells [1]. The majority of secreted proteins in eukaryotes go along the endoplasmic reticulum (ER)-Golgi pathway [2]. This pathway is guided via a signal peptide present on the N-terminus of the protein, also known as the leader sequence. It helps deliver the nascent proteins from ER to the Golgi apparatus, which are then transported to the cellular surface via vesicles [3]. Apart from the classical pathway, i.e., ER-Golgi pathway, some proteins are also secreted through unconventional pathways that are able to secrete the leaderless proteins. Unconventional pathways involve both non-vesicular and vesicular transport. In non-vesicular transport, proteins are secreted into the extracellular space, whereas in vesicular transport, proteins are secreted via vesicles. These vesicular structures comprise a of a variety of classes, and among these classes, exosomes stand out [4,5].

Exosomes belong to a class of extracellular vesicles with endosomal origin, are derived from cells, and range from size 30 to 150 nm [6]. They facilitate interactions with the cellular environment and are extensively found in bodily fluids like urine, saliva, blood, cerebrospinal fluid, bile, breast milk, amniotic fluid, semen, epididymal fluid, and sputum [7]. They are produced in the cytosol as a result of inward budding on late endosomes to form intraluminal vesicles (ILVs) inside a large multivesicular body (MVB) [8]. When MVB merges with the plasma membrane, ILVs are secreted as exosomes into the extracellular environment [9]. Exosomes encompass a compound cargo of contents arising from the original cell, including lipids, DNA, proteins, miRNA, and mRNA (Figure 1) [10]. The content carried by exosomes can change in diseased conditions making it a useful entity for biomarker detection [11]. Exosome-based diagnostics are more specific and sensitive than liquid biopsy or conventional biopsy biomarkers due to their high stability in body fluids [12,13]. In addition, exosomal markers are readily available from most biofluids which makes exosome-based diagnostics labour and cost-effective [14,15]. Since proteins and peptides are the most widely studied macromolecules as biomarkers, identifying and annotating exosomal proteins can help develop the least-invasive novel diagnostic methods as well as therapies for various diseases. The proteins extracted from the circulating exosomes can give us comprehensive information about a specific disease – for example – exosomal proteins can give us important evidence about distal tumors, which is otherwise difficult to obtain due to complex diagnostic methods like tissue biopsy [16]. Extracting proteins from exosomes is more efficient than extracting them from blood, as blood has many substances [17].

**Figure 1:**
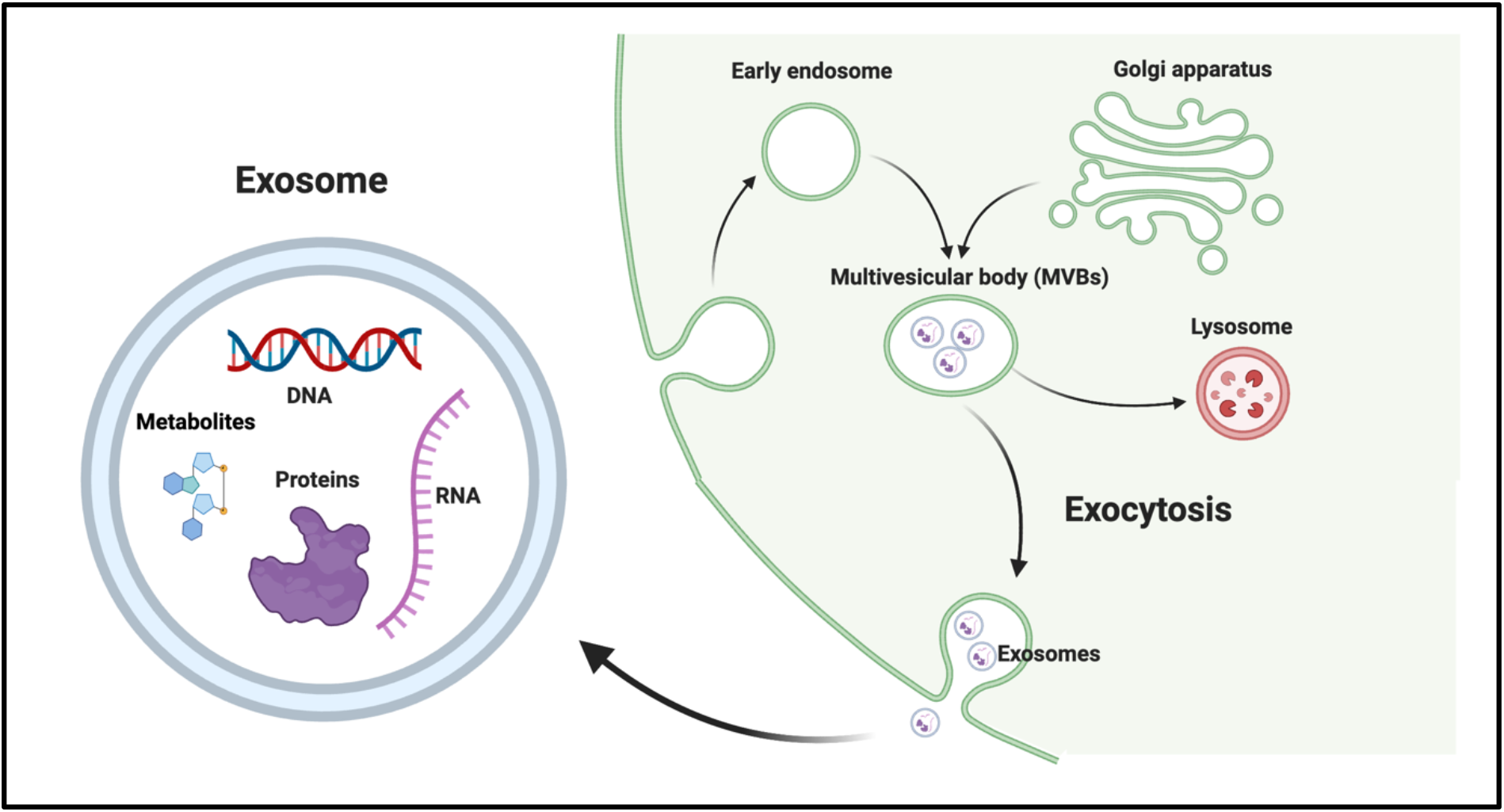
Mechanism of formation of exosomes

Identifying proteins secreted by cells via exosomes has its own challenges, as cells produce wide range of highly similar proteins. In addition, exosomes arise from a range of different cell types and it would be difficult to determine their origin tissue unless they carry extremely specific cargos [18]. Thus, it is crucial to develop a computational method that can predict proteins secreted by exosomes. In this direction, there are several existing methods to predict classical and non-classical secreted proteins that include SRTpred, OutCyte, SecretP, SPRED, and SecretomeP 2.0 [19–23]. None of them has been specifically trained on proteins secreted by exosomes. There is only one method ExoPred that is trained on exosomal proteins for vertebrates [24]. To complement presently available methods, we made a systematic attempt to build a classifier that can annotate human exosomal proteins with high precision. We tried a wide range of model-building techniques, different types of protein features, and a motif-based approach (see Figure 2).

**Figure 2:**
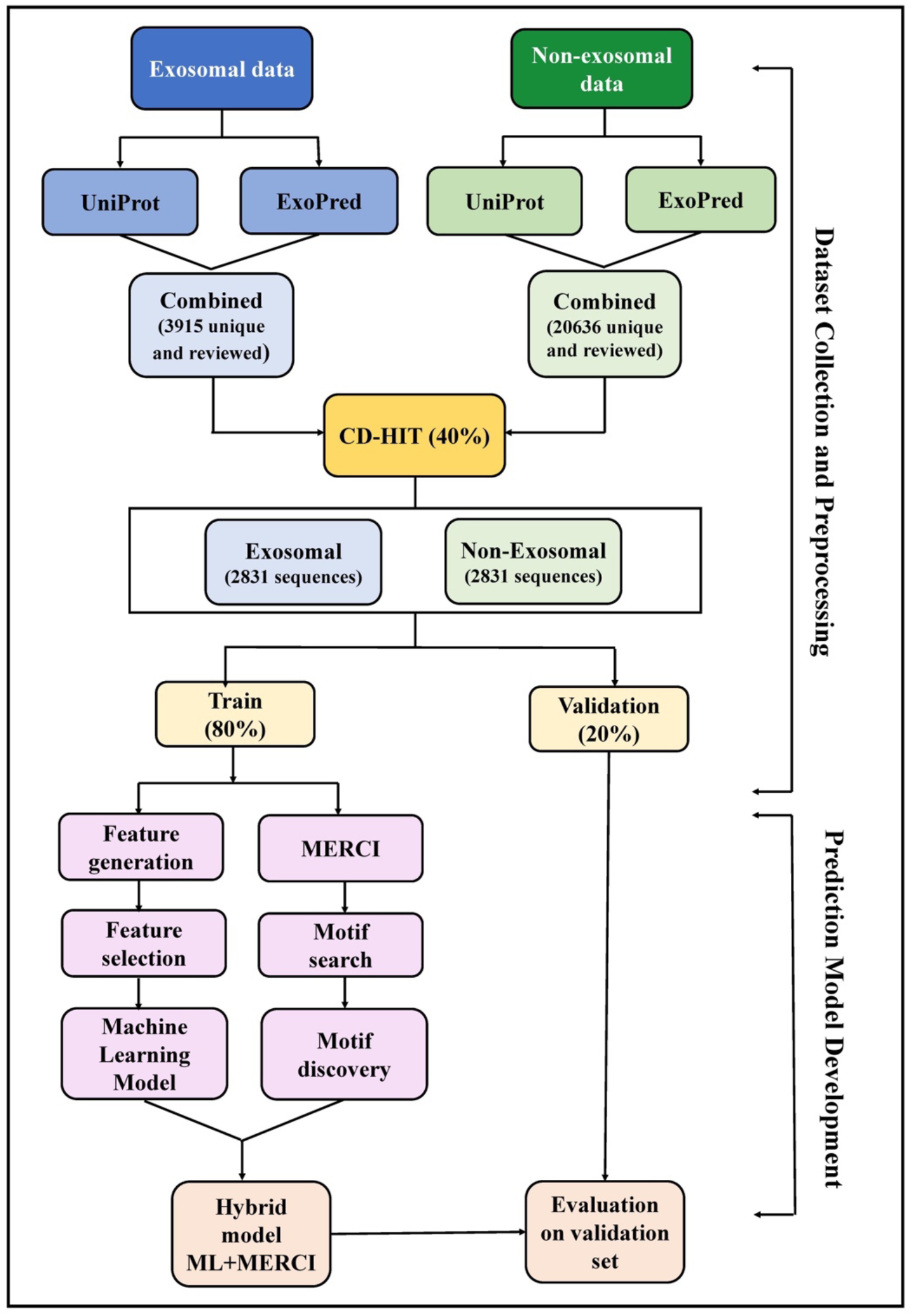
Flowchart of the methodology followed in the study

## Materials and methods

### Compilation and processing of the dataset

The data used in this research work was retrieved from UniProt release 2022_02 (Released on 25 May 2022) and from the ExoPred dataset [24,25]. We retrieved 2178 exosomal proteins from UniProt using the following queries; i) (go:0070062) AND (reviewed:true) AND (organism_id:9606), ii) “extracellular exosome” AND (reviewed:true) AND (organism_id:9606), and iii) “exosome” AND (reviewed:true) AND (organism_id:9606). In addition, we retrieved 2551 exosomal proteins from ExoPred dataset, which are reviewed proteins belonging to human. After compiling the data extracted from UniProt and ExoPred, we had a total of 3915 exosomal proteins. Similarly, we extracted 18207 non-exosomal proteins from UniProt using the following query, NOT (go:0070062) NOT Exosomes NOT “Extracellular exosome” NOT Exosome “ AND (reviewed:true) AND (organism_id:9606). We also combined these non-exosomal proteins with the non-exosomal proteins from ExoPred dataset. Finally, we got 20330 unique non-exosomal proteins after removing duplicates. We also removed proteins consisting of non-standard amino acids “BJOUXZ” and sequences with lengths <55 and >1500. Finally, we obtained 2831 non-redundant exosomal proteins after discarding redundant sequences using CD-HIT software where no two proteins have more than 40% similarity [26]. Similarly, we obtained 10680 non-exosomal proteins after removing redundant sequences. The final dataset contains 2831 exosomal and 2831 non-exosomal (randomly selected from 10680 non-exosomal sequences) proteins.

### Feature generation

To develop a prediction model to classify proteins, we need a set of features for every protein.

A number of feature encoding techniques have been used in previous studies [27–30]. We used a standalone tool called Pfeature to compute numerous features for the proteins, including evolutionary information-based features and composition-based features [31].

#### Composition-based features

The composition-based feature module available on Pfeature provides a vector of 9163 features for every protein in the positive (exosomal) and negative (non-exosomal) dataset like Amino Acid Composition (AAC), Tri-peptide Composition (TPC), Di-peptide Composition (DPC), and many more.

#### Evolutionary features

The evolutionary features of a protein are known to provide additional important information about proteins than its other primary sequence features [32,33]. The evolutionary information can be retrieved by calculating the position-specific scoring matrix (PSSM) profile using Position-Specific Iterated BLAST (PSI-BLAST) for each protein [34]. We have obtained a PSSM-400 composition profile as evolutionary features, which have been described in earlier studies [33]. PSSM-400 is a 20 × 20 dimension vector for a protein sequence which comprises the measure of occurrences of 20 amino acids in the sequence. We have created a PSSM matrix for each sequence which was first normalized within the range of 0 to 1 and converted into a PSSM composition of size 20 × 20 vector using Pfeature software [31].

### Feature selection

It has been shown earlier that all the features extracted from a protein are not relevant, and there is a need to select only the useful ones from a big set of features [35]. To achieve the same, we applied the Recursive Feature Elimination (RFE) feature selection technique using the Scikit-learn package in Python programming language using Logistic Regression as the estimator [36]. We selected the top 20 and top 50 most relevant compositional features and evolutionary features (PSSM composition), respectively. This feature selection method keeps removing the weakest features from the set until a specified number of features has been reached. The features were selected from the standardized data that was obtained using StandardScaler in the Scikit-learn package [37]. The features that were top-ranked were then used to create several machine-learning prediction models for the dataset. The features used in the machine learning (ML) models are described in Supplementary Table S1.

### Similarity search using BLAST

Basic Local Alignment Search Tool (BLAST) version-2.2.29+ is widely used to identify and annotate protein and nucleotide sequences [38]. In this research study, we tried to use BLAST for the identification of exosomal proteins. It is based on the protein sequence similarity with exosomal and non-exosomal protein sequences. The protein query sequences were made to hit against a database of exosomal and non-exosomal protein sequences.

We applied three approaches to identify exosomal sequences, which involved taking into account the top hit, top three hits, and top five hits at various E-value cut-offs. In the first strategy, i.e., first hit, - the sequence is identified as exosomal or non-exosomal based on its first hit against the whole database. However, for the top three and five hits, a voting approach is considered, and a sequence is identified as exosomal if top three or five hits have the maximum of exosomal proteins. The non-exosomal proteins are also characterized in the same manner. For this, a minimum of three or five hits must be available for voting. The performance of these three strategies was recorded for different E-values. Several research have used this methodology to identify a protein sequence [27,33].

### Motif search

It is important to recognize the functional motifs present in the protein or peptide sequences for their functional annotation as well as to classify the negative and positive datasets. In this study, we used Motif Emerging with Classes Identification (MERCI) program to find motifs in both exosomal and non-exosomal protein sequences [39]. MERCI selects specific motifs in the positive dataset by comparing negative and positive input sequences. Hence, to retrieve the specific motifs in exosomal and non-exosomal protein sequences, we followed a two-step procedure that involved - a) Providing exosomal proteins as positive input and non-exosomal proteins as negative input and finding motifs for exosomal protein sequences, b) Reversing the order for positive and negative input to find motifs for non-exosomal protein sequences.

We used different options available in MERCI to extract motifs that are exclusive as well as inclusive to both sets. By default, MERCI takes the maximal frequency of the negative sequences (fn) as zero, which gives only exclusive motifs, i.e., the motifs that are not common in positive and negative sets. We increased this value to fn = 8 to obtain inclusive motifs as well. Within the exclusive and inclusive motifs, we obtained a different kinds of motifs by specifying some values that include – a) No gap, b) Gap = 1, c) Gap = 2, and d) Class = Koolman-Rohm. After that, the unique proteins containing motifs were selected to compute the overall coverage of motifs in the protein sequences.

### Machine Learning (ML) classifiers

We have employed several ML algorithms to differentiate between exosomal and non-exosomal proteins. These algorithms involve Gaussian Naïve Bayes (GNB), K-Nearest Neighbours (KNN), Decision Tree (DT), Extreme Gradient Boosting (XGB), Logistic Regression (LR), Support Vector classifier (SVC), and Random Forest (RF). The parameters of these algorithms were optimized using hyperparameter tuning.

### Performance metrics calculation and cross-validation

The whole dataset was divided into the ratio of 80:20, where 80% comprised the training and 20% validation data. The five-fold cross-validation technique was applied to the 80% of the training data to assess the ML models, and the remaining 20% was kept unknown to the models. In five-fold cross-validation technique, the 80% of training data is split into five parts where four folds are used for training and the left one fold is used as a test set for internal validation purposes. This procedure is reiterated five times so that every fold gets a chance to be the test fold. The ML models used in this study have been evaluated using performance metrics which include parameters dependent and independent of the threshold. The different standard evaluation metrics that have been used in this study include sensitivity, specificity, Matthews correlation coefficient (MCC), accuracy, and Area under the Receiver Operating Characteristics (AUROC). Out of these, AUROC is threshold-independent, and the rest of the parameters are threshold-dependent. These metrics have been previously used in studies to estimate the performance of ML models [40,41].

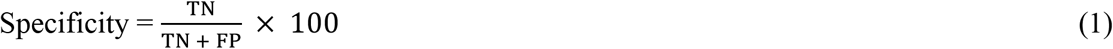

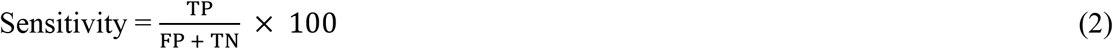

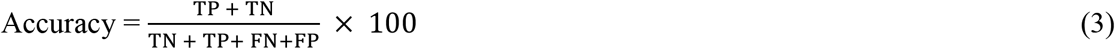

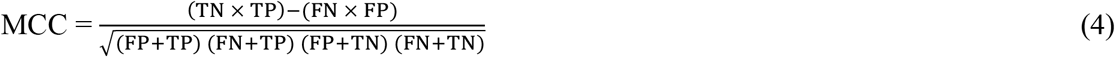

Where, TP, FP, TN, and FN are true positive, false positive, true negative, and false negative, respectively.

### Hybrid model

To improve the prediction of ML models, we applied a hybrid approach that integrates and employs the various results obtained in this study. The hybrid approach uses a weighted scoring method in which the scores are calculated by combining two methods (i) Motif-based approach and (ii) ML-based methods. In this hybrid model, we assigned a weight of +0.5 if a protein sequence had an exosomal motif and −0.5 if it had a non-exosomal motif, and 0 if no motif was found. These scores were combined with the ML scores forming an overall score. The sequences were categorized as exosomal and non-exosomal by analyzing the overall scores at a range of threshold values. A number of studies have used this hybrid approach earlier [27,42].

## Results

### Amino acid composition analysis

After analysing and comparing the amino acid composition of exosomal and non-exosomal proteins, we discovered that exosomal sequences have a comparatively higher amount of aspartic acid, glutamic acid, isoleucine, lysine, asparagine, valine, tyrosine, and lower amounts of cysteine, histidine, leucine, proline, arginine, serine, and tryptophan. The graph of average amino acid composition comparison between exosomal and non-exosomal protein classes is illustrated in Figure 3.

**Figure 3:**
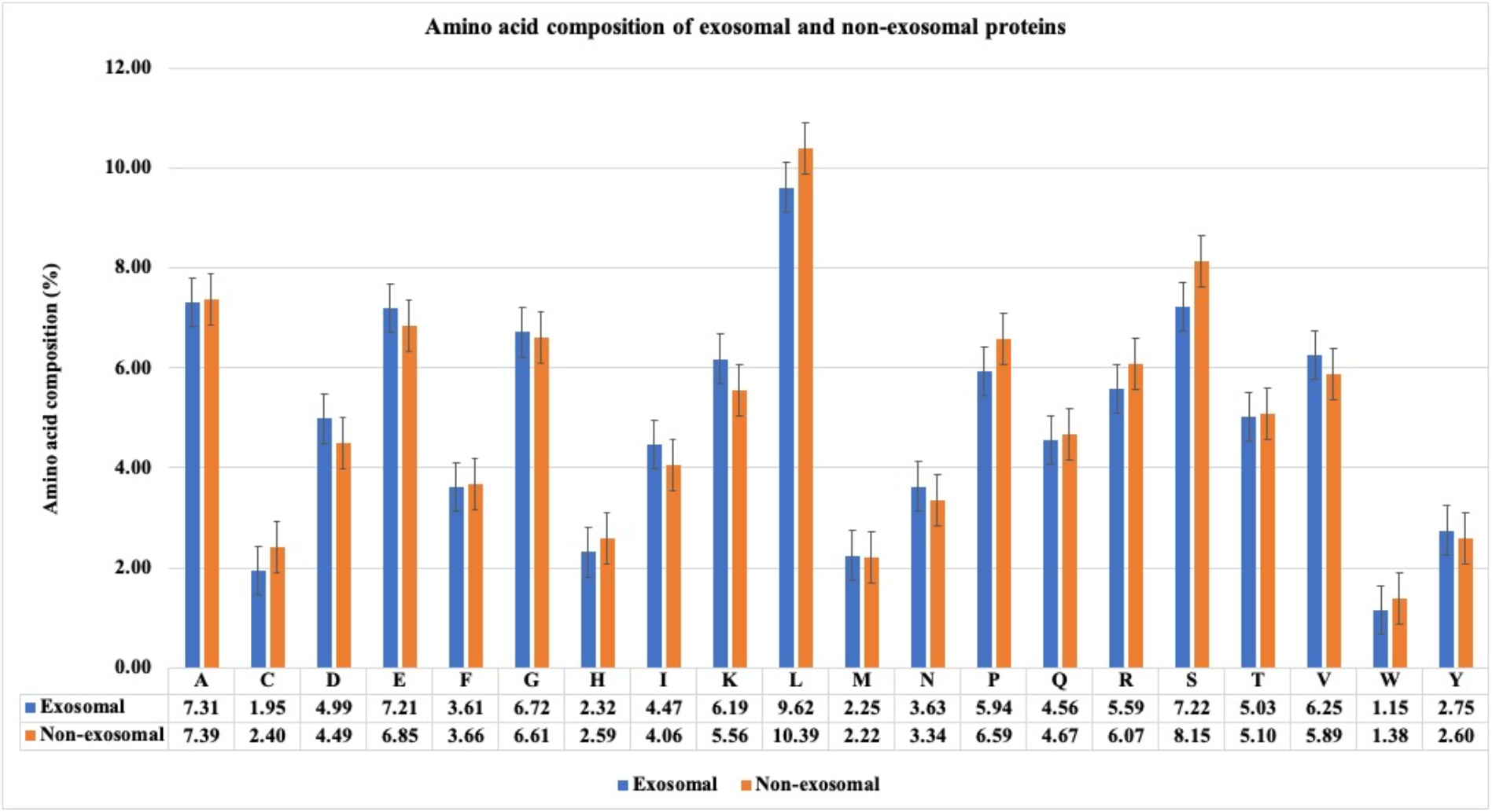
Amino acid composition analysis for exosomal and non-exosomal proteins

### BLAST performance

BLAST is widely used to annotate and recognize the role of a query protein sequence on the basis of similarity search. We attempted to utilize BLAST in this study to classify proteins as exosomal and non-exosomal. We used five-fold cross-validation to evaluate the performance of BLAST. Firstly, the sequences in four folds are used to create a database, and the sequences present in the fifth fold are used to hit against the sequences present in respective database. This procedure was repeated five times. To evaluate BLAST performance on the validation dataset, we constructed a database using all the training sequences, and those in the validation set were searched against the database. We used BLAST in three ways – a) Top hit, b) Top 3 hits, and c) Top 5 hits. The top hit is a common method that assigns a class to the protein based on the first hit, whereas, top 3 and top 5 criteria assign a class to the protein on the basis of the class that is appearing the maximum number of times in the first 3 and 5 hits. We have considered top 3 and 5 hits as sometimes the top hit is not always the most relevant one. However, even after trying all these criteria, we were getting a large number of false positives and false negatives. We obtained 18.06%, 11.08%, and 8.39% sensitivity (number of correct hits) for the training dataset, whereas the sensitivity of 17.92%, 11.39%, and 8.65% was obtained for the validation dataset, for top 1, top 3, and top 5 hits respectively. With the increment of the E-value the error rate was also increasing. For the training set, we got 9.45%, 4.75%, and 3.2%, and for the validation set, we got 12.97%, 7.41%, and 6% for top 1, top 3, and top 5 hits respectively. The results for BLAST are shown in Table 1.

**Table 1:**
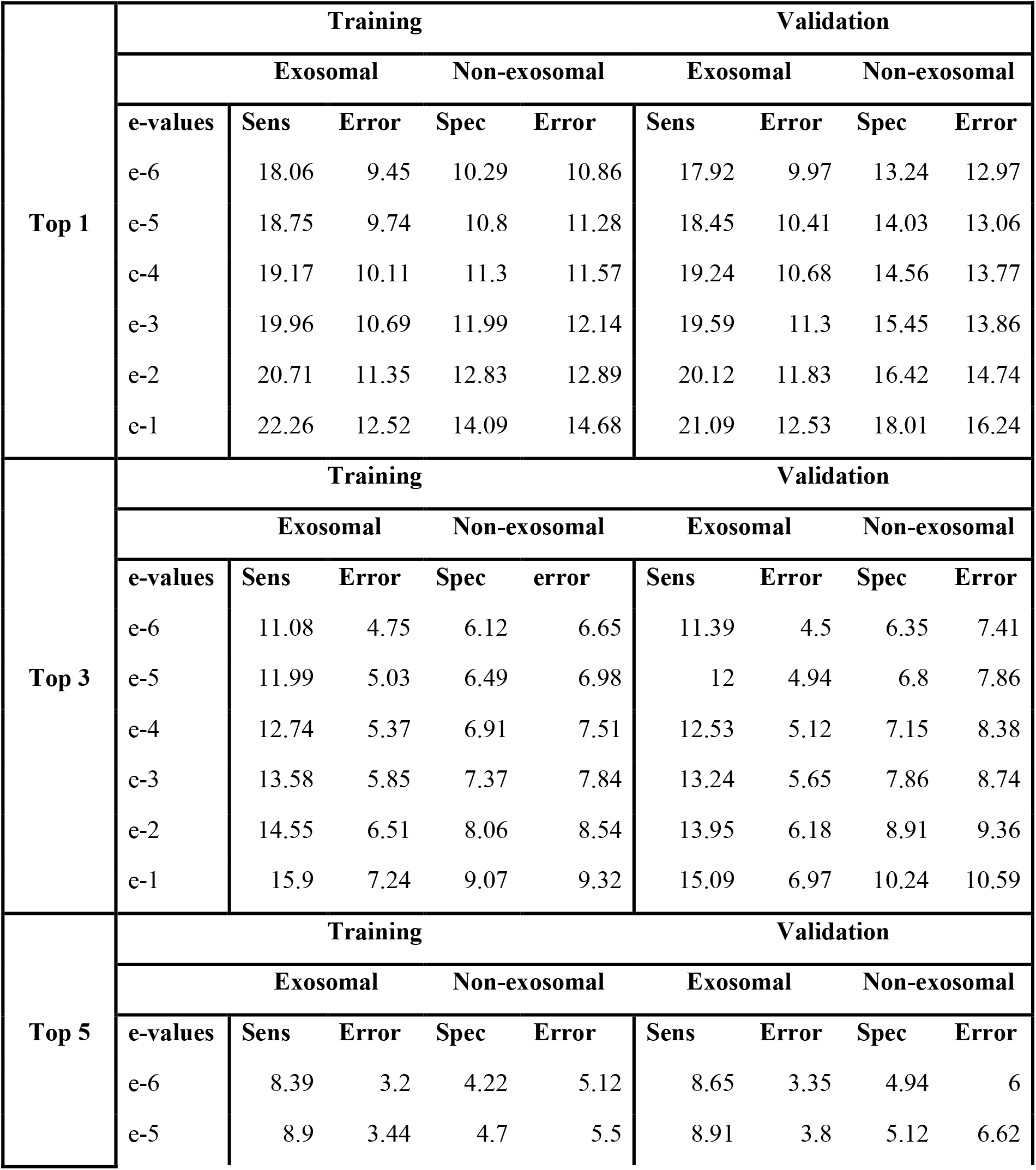
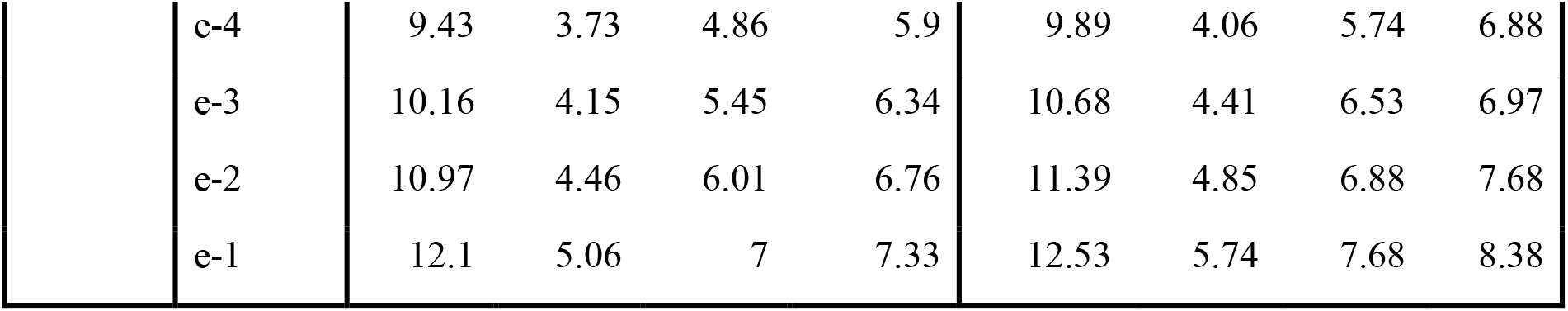
Results for top 1, top 3, and top 5 hits in BLAST for validation set searched against training set database (sens = sensitivity, spec = specificity)

### Machine learning models

For each protein sequence in the dataset, a total of 9163 features were computed that constituted more than ten types of compositional features. Along with the composition-based features, evolutionary features were also computed using Pfeature software. The relevant features were then selected using the feature selection technique - RFE and the ML models were developed using the scikit-learn package in Python [37].

### Compositional features

The amino acid composition (AAC) for the exosomal and non-exosomal proteins was calculated to build the ML models. The random forest ML model was observed to perform well than other models and was able to achieve an AUROC of 0.71 and 0.70 on training and validation sets, respectively. The performance of AAC for the dataset is given in Table 2.

**Table 2:** Results for Machine Learning (ML) models developed for Amino Acid Composition (AAC) and Position-Specific Scoring Matrix (PSSM) composition features

| Amino Acid Composition (AAC) |  |  |  |  |  |  |  |  |  |  |
| --- | --- | --- | --- | --- | --- | --- | --- | --- | --- | --- |
| Training |  |  |  |  |  | Validation |  |  |  |  |
| Model | Sens | Spec | Acc | AUC | MCC | Sens | Spec | Acc | AUC | MCC |
| DT | 59.75 | 59.92 | 59.84 | 0.62 | 0.20 | 50.18 | 61.75 | 56.13 | 0.58 | 0.12 |
| **RF** | **64.58** | **66.15** | **65.36** | **0.71** | **0.31** | **64.91** | **65.01** | **64.96** | **0.70** | **0.30** |
| **LR** | 63.04 | 65.21 | 64.12 | 0.69 | 0.28 | 59.09 | 63.47 | 61.34 | 0.67 | 0.23 |
| **XGB** | 63.92 | 64.68 | 64.30 | 0.70 | 0.29 | 61.64 | 65.35 | 63.55 | 0.70 | 0.27 |
| **KNN** | 62.87 | 65.30 | 64.08 | 0.69 | 0.28 | 64.36 | 60.89 | 62.58 | 0.68 | 0.25 |
| **GNB** | 61.68 | 62.32 | 62.00 | 0.67 | 0.24 | 60.00 | 62.26 | 61.17 | 0.64 | 0.22 |
| **SVC** | 64.93 | 64.99 | 64.96 | 0.70 | 0.30 | 62.36 | 61.92 | 62.14 | 0.68 | 0.24 |
| **PSSM Composition** |  |  |  |  |  |  |  |  |  |  |
| **Training** |  |  |  |  |  | **Validation** |  |  |  |  |
| **Model** | **Sens** | **Spec** | **Acc** | **AUC** | **MCC** | **Sens** | **Spec** | **Acc** | **AUC** | **MCC** |
| **DT** | 57.87 | 58.26 | 58.06 | 0.62 | 0.16 | 56.36 | 54.72 | 55.52 | 0.59 | 0.11 |
| **RF** | 68.30 | 66.76 | 67.54 | 0.73 | 0.35 | 66.73 | 65.87 | 66.28 | 0.71 | 0.33 |
| **LR** | **67.91** | **67.29** | **67.60** | **0.73** | **0.35** | **65.09** | **67.24** | **66.20** | **0.72** | **0.32** |
| **XGB** | 65.28 | 64.80 | 65.04 | 0.71 | 0.30 | 64.91 | 61.41 | 63.11 | 0.70 | 0.26 |
| **KN** | 66.37 | 64.93 | 65.66 | 0.71 | 0.31 | 65.46 | 59.18 | 62.22 | 0.69 | 0.25 |
| **GNB** | 67.51 | 60.48 | 64.02 | 0.68 | 0.28 | 63.82 | 59.52 | 61.61 | 0.67 | 0.23 |
| **SVC** | 67.82 | 66.89 | 67.36 | 0.73 | 0.35 | 65.82 | 65.87 | 65.84 | 0.71 | 0.32 |

### Evolutionary features

The ML models were also developed on the basis of evolutionary information. To obtain the evolutionary information, we computed the PSSM profiles of each protein which were then fed to our ML models. It was observed that the logistic regression model was performing best on these features and achieved the AUROC of about 0.73 and 0.72 for training and validation sets. The performance of PSSM-based models is given in Table 2.

### Feature selection

A total of 9163 composition-based features were generated using Pfeature software, which were short-listed to top 20 features using the RFE feature selection method based on the logistic regression estimator. These 20 features attained an AUROC of 0.71 on training and 0.71 on validation sets based on the support vector classifier (SVC). We also selected the top 50 features for evolutionary information-based features using the same technique, which yielded the AUROCs of 0.74 on training and 0.71 on validation sets using a random forest classifier. We also compiled the top 20 compositional and top 50 evolutionary features, which resulted in a matrix of a total of 70 features. The combination of these features was able to achieve the AUROCs of 0.75 on training and 0.73 on validation sets. The results for all the selected features are shown in Table 3.

**Table 3:**
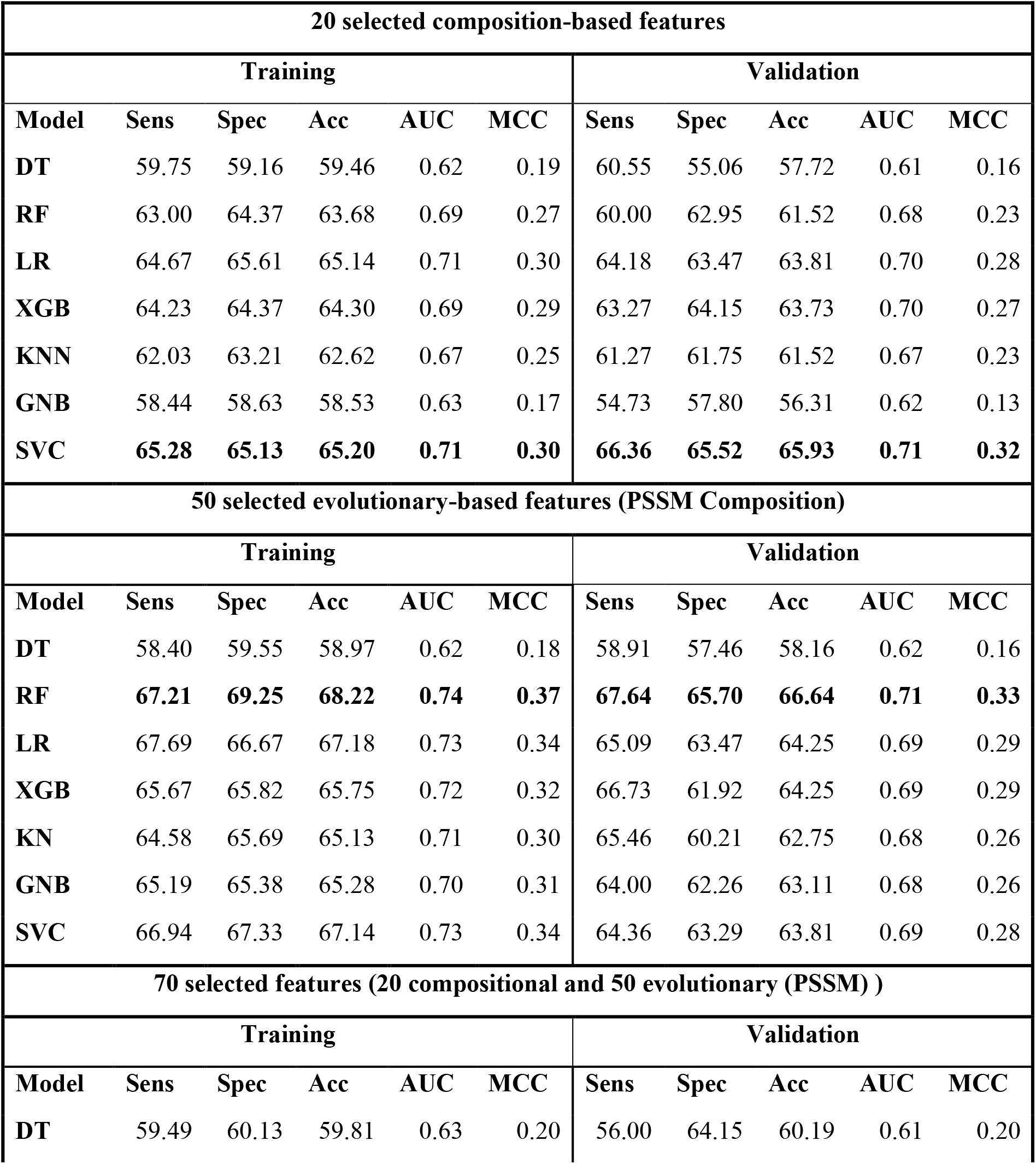
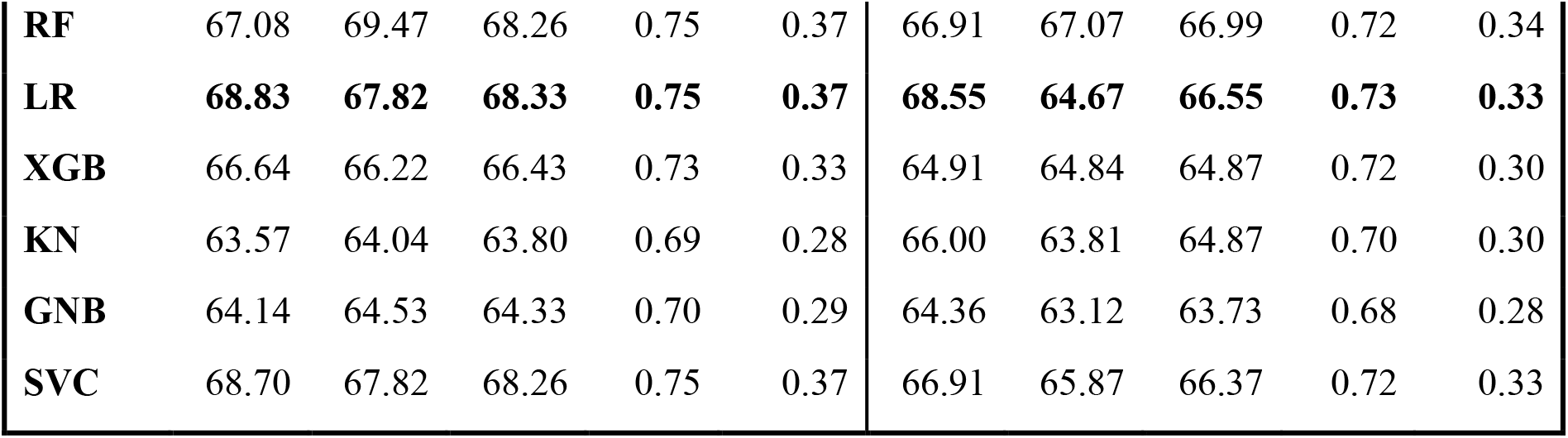
Results for Machine Learning (ML) models developed for top 20 composition features, top 50 evolutionary (PSSM) features, and combination of top selected composition and evolutionary (PSSM) features

### Top selected features

The top compositional features include Amino Acid Index (AAI), Atom Composition (ATC), Pseudo Amino Acid Composition (PAAC), Shannon Entropy (SEP), Quasi-Sequence Order (QSO), and Shannon Entropy of Residue Level (SER). Amongst these, it was observed that three of the relevant features include SER, QSO, and PAAC of tryptophan (W) which indicates that tryptophan can be a significant amino acid for differentiating between exosomal and non-exosomal proteins. It is found to be present in lower concentration in exosomal proteins in the amino-acid composition analysis. Along with this, it was observed that atom composition of Nitrogen and Sulphur were also two of the critical features in predicting exosomal proteins.

### Motif search

We attempted to identify the exclusive and inclusive set of motifs present in exosomal and non-exosomal proteins using the publicly available MERCI program. To achieve this, we extracted motifs using different parameters like – a) No gap, b) Gap = 1, c) Gap = 2, and d) Class = Koolman-Rohm. By default, MERCI takes fn (maximal frequency in negative sequences) as zero, which gives exclusive motifs; we increased it to fn = 8 to get inclusive motifs for both negative and positive datasets. Altogether, MERCI provided 89 motifs in exosomal and 130 motifs in a non-exosomal set that covered 1441 exosomal and 1373 non-exosomal sequences. The top 5 motifs in each category for the exclusive and inclusive sets and the number of sequences they occurred in is given in Table 4. It is observed that most of the motifs crucial for predicting exosomal proteins consisted of aliphatic amino acids except tyrosine (Y). Aromatic amino acids like tryptophan (W) and phenylalanine (F) were not found in any of the important motifs.

**Table 4:**
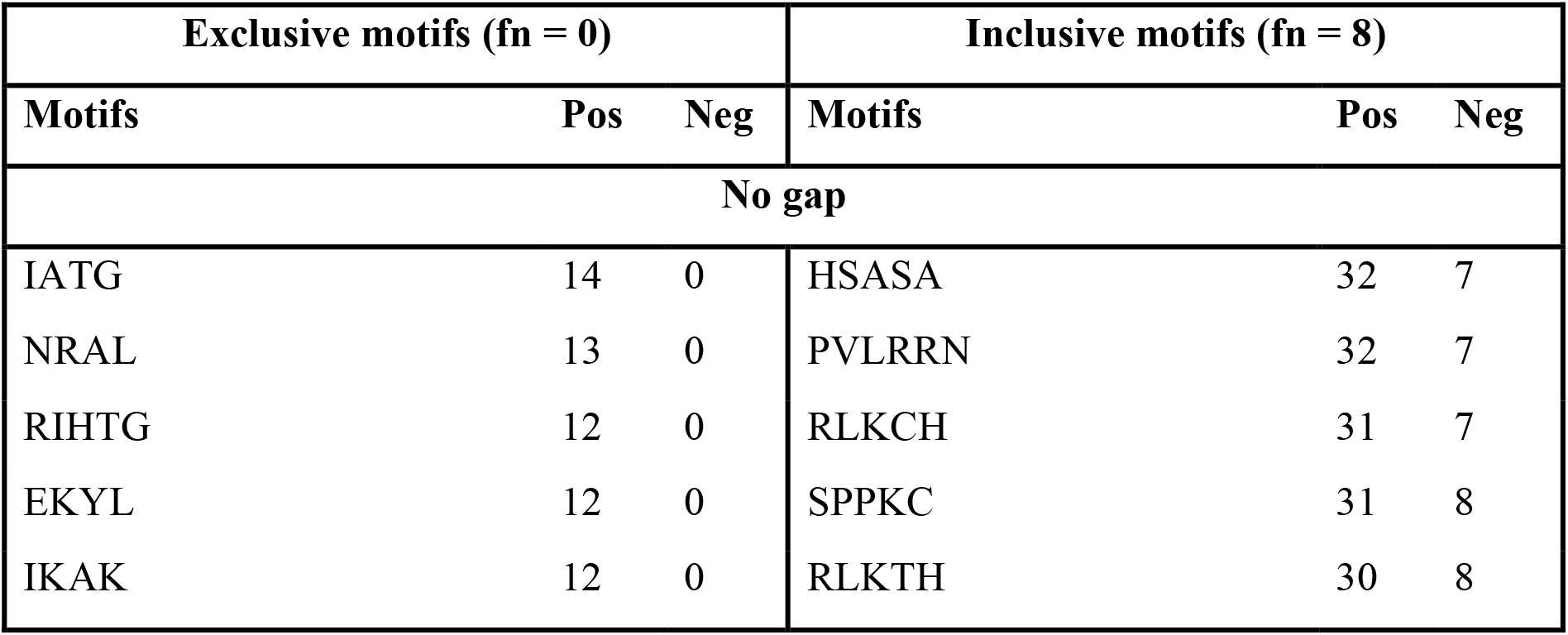
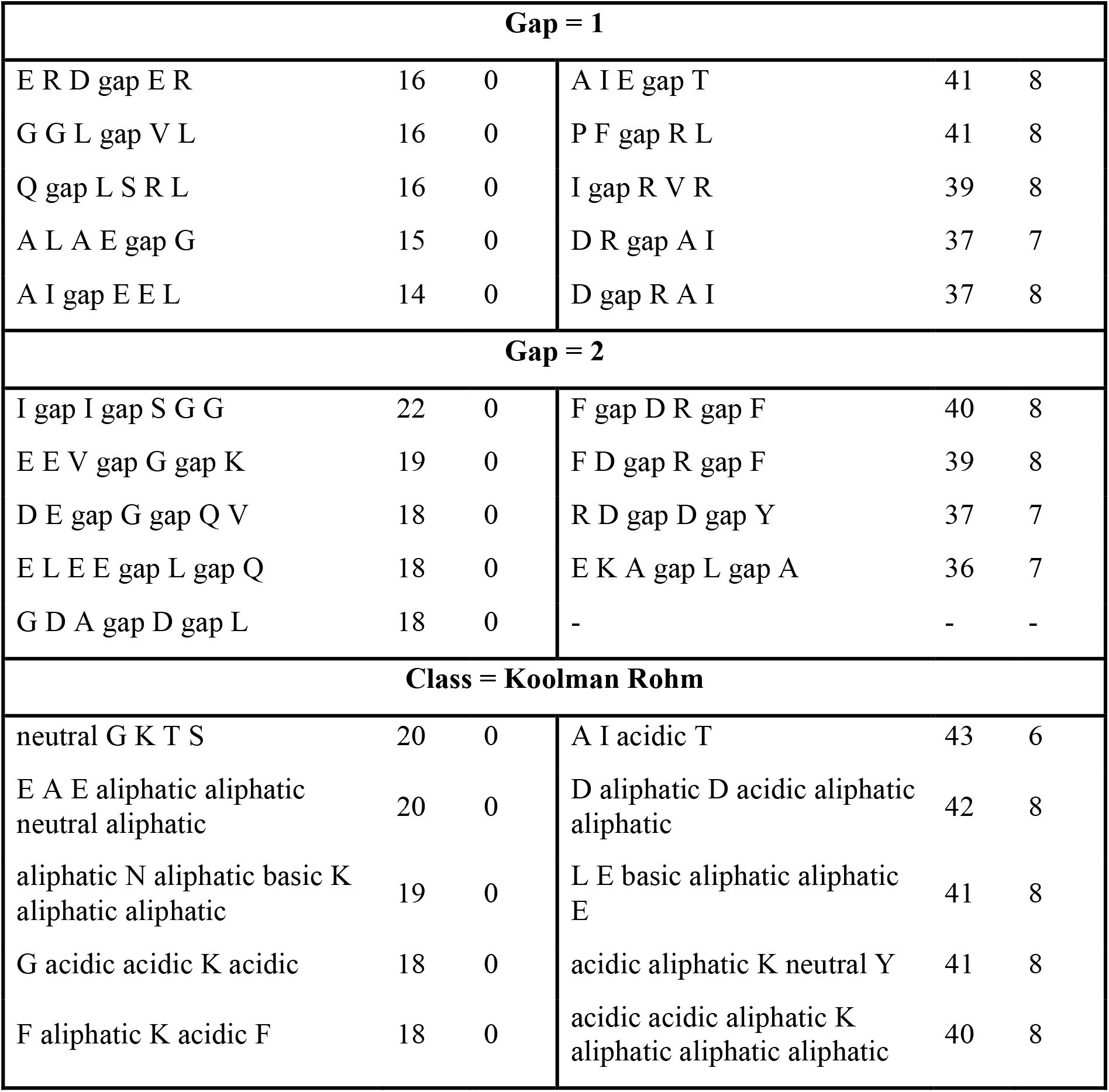
Top 5 motifs exclusive and inclusive motifs and the number of sequences in which they occurred a) No gap, b) Gap=1, c) Gap=2, and d) Class=Koolman Rohm (fn = maximal frequency in negative sequences, pos = occurrence in positive sequences, neg = occurrence in negative sequences)

### Hybrid approach

Since we were getting a good sequence coverage using the motif search, we decided to combine motif prediction with ML-based prediction. In this hybrid model, we allotted a protein sequence a score of +0.5 if an exosomal motif was present, −0.5 if a non-exosomal motif was present, and 0 if none were present. These scores were compiled with the scores obtained from ML-based predictions. After merging the motif prediction scores with random forest model prediction based on AAC, we acquired an AUROC of 0.86 for training and 0.84 for the validation dataset. We also merged the motif prediction scores with Random Forest model prediction based on the top 70 features (20 compositional and 50 evolutionary), which attained an AUROC of 0.82 for training and 0.85 for the validation set. The performance of other models has been explained in Table 5. The AUROC plots for training and validation sets for ML and hybrid models are illustrated in figure 4.

**Table 5:** Results of Hybrid approach (a) MERCI+ ML (AAC) (b) MERCI +ML (top 70 features)

| AAC |  |  |  |  |  |  |  |  |  |  |
| --- | --- | --- | --- | --- | --- | --- | --- | --- | --- | --- |
| Training |  |  |  |  |  | Validation |  |  |  |  |
| Model | Sens | Spec | Acc | AUC | MCC | Sens | Spec | Acc | AUC | MCC |
| DT | 72.42 | 73.71 | 73.06 | 0.8 | 0.46 | 66.73 | 74.44 | 70.7 | 0.78 | 0.41 |
| RF | **76.85** | **77.31** | **77.08** | **0.86** | **0.54** | **77.27** | **75.64** | **76.43** | **0.84** | **0.53** |
| LR | 76.46 | 75.89 | 76.18 | 0.85 | 0.52 | 74.91 | 74.44 | 74.67 | 0.84 | 0.49 |
| XGB | 76.11 | 76.69 | 76.4 | 0.85 | 0.53 | 76.55 | 75.99 | 76.26 | 0.84 | 0.53 |
| KNN | 76.33 | 75.93 | 76.13 | 0.85 | 0.52 | 78.18 | 72.9 | 75.46 | 0.84 | 0.51 |
| GNB | 73.91 | 73.67 | 73.79 | 0.82 | 0.48 | 73.27 | 70.84 | 72.02 | 0.79 | 0.44 |
| SVC | 76.24 | 77.58 | 76.9 | 0.85 | 0.54 | 74.91 | 74.44 | 74.67 | 0.84 | 0.49 |
| 70 selected features (20 compositional and 50 evolutionary (PSSM) ) |  |  |  |  |  |  |  |  |  |  |
| Training |  |  |  |  |  | Validation |  |  |  |  |
| Model | Sens | Spec | Acc | AUC | MCC | Sens | Spec | Acc | AUC | MCC |
| DT | 69.31 | 69.51 | 69.41 | 0.76 | 0.39 | 72.00 | 72.56 | 72.29 | 0.79 | 0.45 |
| RF | **74.62** | **74.99** | **74.80** | **0.82** | **0.50** | **79.82** | **76.16** | **77.93** | **0.85** | **0.56** |
| LR | 74.53 | 75.03 | 74.78 | 0.82 | 0.50 | 77.45 | 76.16 | 76.79 | 0.85 | 0.54 |
| XGB | 73.87 | 73.39 | 73.63 | 0.82 | 0.47 | 75.82 | 75.30 | 75.55 | 0.85 | 0.51 |
| KNN | 72.95 | 71.74 | 72.35 | 0.79 | 0.45 | 78.36 | 74.27 | 76.26 | 0.84 | 0.53 |
| GNB | 71.59 | 71.43 | 71.51 | 0.79 | 0.43 | 74.36 | 71.18 | 72.73 | 0.81 | 0.46 |
| SVC | 75.27 | 74.81 | 75.04 | 0.82 | 0.50 | 78.00 | 75.99 | 76.96 | 0.85 | 0.54 |

**Figure 4:**
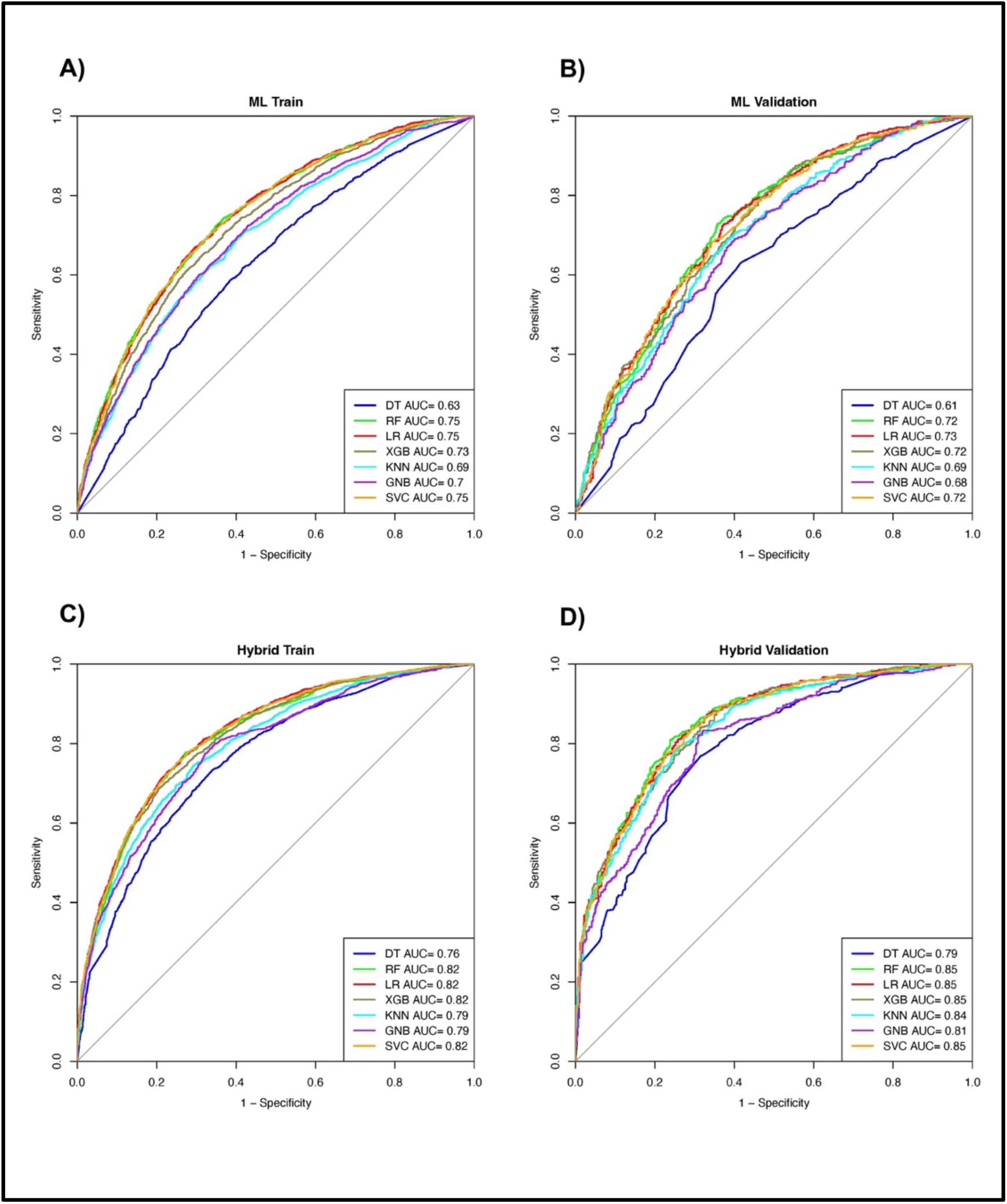
AUROC plots for A) Training set in ML model, B) Validation set in ML model, C) Training set in the hybrid model, and D) Validation set in the hybrid model

### Web server development

We developed a web server – ExoProPred (https://webs.iiitd.edu.in/raghava/exopropred/) to predict exosomal proteins. We have integrated our two best-performing hybrid models – a) AAC combined with MERCI and b) Top 70 features combined with MERCI. The web server incorporates the key modules, including a) prediction, b) motif scan, and c) download. The “prediction module” allows users to submit their query protein sequences in FASTA format. This module can predict exosomal and non-exosomal proteins effectively. The second module – “motif scan,” can identify the motifs present in exosomal and non-exosomal protein sequences using the MERCI software. This module can also scan or map the motifs present in the protein sequence query entered by the user and differentiates between them as exosomal or non-exosomal sequences. The web server has been developed on a responsive HTML template and is compatible with various operating systems. We also built a python-based standalone package of ExoProPred to help users easily predict and classify the sequences at a large scale, and can be downloaded from the “download module” on the web server.

### Comparison with other prediction tools

Presently, there is only one tool that predicts exosomal proteins – ExoPred. The other tools, such as SecretomeP 2.0, and Outcyte, predict whether the protein is following an unconventional pathway or not [19,20]. We entered our validation set of 569 sequences into each of the servers after subtracting the sequences taken from the ExoPred dataset, and performed a comparative analysis. ExoPred was able to predict the sequences with 66.08% accuracy [24]. However, it had very low sensitivity but high specificity, which means it is able to predict the non-exosomal sequences but not able to classify exosomal sequences correctly. For SecretomeP 2.0, we selected the “mammalian” option on the web server, and as indicated on their webpage, proteins with “NN score” higher than 0.6 are said to be secreted via unconventional pathways. After setting this threshold, we found that it was able to predict exosomal sequences with 54.83% accuracy [22]. In Outcyte web server, we selected Outcyte UPS (Unconventional protein secretion option) and obtained an accuracy of 61.16% for our validation set, with a low sensitivity and comparatively higher specificity [19]. On entering these sequences on our web server – ExoProPred, we obtained an accuracy of 79.4%, which is higher than all the above-mentioned tools. ExoProPred is also able to achieve a balanced sensitivity and specificity along with the highest accuracy. The full comparison of prediction by web servers is given in Table 6.

**Table 6:** Comparison of prediction by web servers ExoPred, SecretomeP 2.0, and Outcyte with ExoProPred on validation dataset

| Prediction Model | TP | FP | TN | FN | Sens | Spec | Acc |
| --- | --- | --- | --- | --- | --- | --- | --- |
| **Exopred** | 26 | 51 | 350 | 142 | 15.48% | 87.28% | 66.08% |
| **SecretomeP 2.0** | 80 | 169 | 232 | 88 | 47.62% | 57.85% | 54.83% |
| **Outcyte** | 47 | 100 | 301 | 121 | 27.97% | 75.06% | 61.16% |
| **ExoProPred** | **133** | **83** | **318** | **34** | **79.64%** | **79.30%** | **79.40%** |

## Discussion

There is a need to develop non-invasive diagnostic methods and therapies to prevent patients from going through painful medical procedures to get treatment. Exosomal biomarkers can be found in body fluids (saliva, blood, urine, etc.) in abundance and can be used to detect a disease or develop a treatment for different types of conditions [11]. These biomarkers are derived from the parent cells and are even more specific and sensitive than those extracted directly from the body fluids because the exosome is highly stable and non-immunogenic [43]. Among the disease biomarkers, proteins have been broadly studied and can be used for diagnosis, prognosis, and treatment of specific diseases. However, it is difficult to identify these proteins as they are extremely similar to those produced by the cells, and there is a mixture of exosomes in the biofluids derived from different types of cells [18]. To overcome this limitation, we made an effort to develop a prediction server ExoProPred that classifies the proteins into exosomal and non-exosomal.

In this study, we created a dataset for 2831 exosomal and 2831 non-exosomal proteins extracted from UniProt and ExoPred server [24,25]. A number of features were generated for the protein sequences using the Pfeature tool, and only the relevant features were mined from a big set of features using Recursive Feature Elimination (RFE). Amino Acid Index (AAI), Atom Composition (ATC), Pseudo Amino Acid Composition (PAAC), Shannon Entropy (SEP), Quasi-Sequence Order (QSO), and Shannon Entropy of Residue Level are the top compositional features (SER). The SER, QSO, and PAAC of tryptophan (W) were shown to be three of the relevant properties indicating that tryptophan can be a helpful amino acid for distinguishing between exosomal and non-exosomal proteins. The examination of the amino-acid composition reveals that tryptophan is decreased in exosomal proteins as compared to non-exosomal. Additionally, it was shown that the atom compositions of nitrogen and sulfur were two other crucial characteristics for predicting exosomal proteins.

Besides the development of machine learning (ML) models on selected essential features, we also applied the BLAST tool to identify the exosomal proteins, as this tool has been widely used to annotate the query proteins [38]. However, we were not able to obtain very high performance with BLAST. An explanation for this could be that exosomal proteins are very similar to the proteins present in their parent cell; hence, it must be difficult to point out which protein belongs to the exosome. We decided to exclude BLAST-based performance from our hybrid model, and to boost the performance of the hybrid model; we added a motif-based approach using MERCI in which we obtained 89 exosomal and 130 non-exosomal motifs covering 1441 exosomal and 1373 non-exosomal sequences [39]. On analyzing the top motifs obtained for the classification of exosomal and non-exosomal sequences, we observed that most motifs contained aliphatic amino acids except the presence of tyrosine (Y) at some places. Aromatic amino acids tryptophan (W) and phenylalanine (F) were rarely seen in any of the motifs found crucial for the detection of exosomal sequences. This information complements the observation in the top 20 composition features, which proved that the composition of tryptophan (W) was an essential factor in predicting exosomal sequences. The motif-based approach was able to cover a high amount of exosomal sequences. Hence, we combined this approach with the top-selected features to develop a hybrid model to predict exosomal protein sequences. We are able to achieve an accuracy of 78% and an AUROC of 0.85 with balanced specificity and sensitivity on an independent validation set. In addition to this, we obtained the highest accuracy for a validation set of 569 sequences when compared to other prediction web servers like Outcyte, ExoPred, and SecretomeP 2.0.

We have created a platform that allows users to classify exosomal and non-exosomal protein sequences. We believe that our study will be beneficial for scientists studying protein or peptide diagnostics and therapies. The web server is freely available to help the scientific community and to encourage the use of this prediction method in research. In this web server, we have implemented our best-performing model – the hybrid model. We hope researchers worldwide will make use of our prediction method for the development of more effective and precise protein or peptide-based diagnostic or therapeutic techniques for a range of ailments.

## Author’s contributions

AA, NS, and, NLD collected and processed the data. AA, SP, NS, and NLD implemented the algorithms. AA and SP developed the prediction models. AA and SP developed the front end and back end of the web server. AA, DK, and GPSR prepared the manuscript. GPSR conceived and coordinated the project. All authors have read and approved the final manuscript.

## Acknowledgements

Authors are thankful to Council of Scientific and Industrial Research (CSIR), Department of Biotechnology (DBT), Department of Science and Technology (DST-INSPIRE) and DBT-RA program for providing fellowships and the financial support. Authors are also thankful to the Department of Computational Biology, IIITD, New Delhi for infrastructure and facilities. We thank the Department of Biotechnology (DBT) for providing an infrastructure grant to the institute. We would like to acknowledge that figures were created using BioRender.com.

## Data Availability Statement

All the datasets generated in this study are available at https://webs.iiitd.edu.in/raghava/exopropred/dataset.php.

